# Heart Rate n-Variability (HRnV) and Its Application to Risk Stratification of Chest Pain Patients in the Emergency Department

**DOI:** 10.1101/738989

**Authors:** Nan Liu, Dagang Guo, Zhi Xiong Koh, Andrew Fu Wah Ho, Feng Xie, Takashi Tagami, Jeffrey Tadashi Sakamoto, Pin Pin Pek, Bibhas Chakraborty, Swee Han Lim, Jack Wei Chieh Tan, Marcus Eng Hock Ong

**Author notes:** **Corresponding Author** Nan Liu, Centre for Quantitative Medicine and Programme in Health Services and Systems Research, Duke-NUS Medical School, Academia, 20 College Road, Singapore 169856.

## Abstract

**Background:** Chest pain is one of the most common complaints among patients presenting to the emergency department (ED). Causes of chest pain can be benign or life threatening, making accurate risk stratification a critical issue in the ED. In addition to the use of established clinical scores, prior studies have attempted to create predictive models with heart rate variability (HRV). In this study, we proposed heart rate n-variability (HRnV), an alternative representation of beat-to-beat variation in electrocardiogram (ECG) and investigated its association with major adverse cardiac events (MACE) for ED patients with chest pain.

**Methods:** We conducted a retrospective analysis of data collected from the ED of a tertiary hospital in Singapore between September 2010 and July 2015. Patients >20 years old who presented to the ED with chief complaint of chest pain were conveniently recruited. Five to six-minute single-lead ECGs, demographics, medical history, troponin, and other required variables were collected. We developed the HRnV-Calc software to calculate HRnV parameters. The primary outcome was 30-day MACE, which included all-cause death, acute myocardial infarction, and revascularization. Univariable and multivariable logistic regression analyses were conducted to investigate the association between individual risk factors and the outcome. Receiver operating characteristic (ROC) analysis was performed to compare the HRnV model (based on leave-one-out cross-validation) against other clinical scores in predicting 30-day MACE.

**Results:** A total of 795 patients were included in the analysis, of which 247 (31%) had MACE within 30 days. The MACE group was older and had a higher proportion of male patients. Twenty-one conventional HRV and 115 HRnV parameters were calculated. In univariable analysis, eleven HRV parameters and 48 HRnV parameters were significantly associated with 30-day MACE. The multivariable stepwise logistic regression identified 16 predictors that were strongly associated with the MACE outcome; these predictors consisted of one HRV, seven HRnV parameters, troponin, ST segment changes, and several other factors. The HRnV model outperformed several clinical scores in the ROC analysis.

**Conclusions:** The novel HRnV representation demonstrated its value of augmenting HRV and traditional risk factors in designing a robust risk stratification tool for patients with chest pain at the ED.

## Background

Chest pain is one of the most common presenting complaints in the emergency department (ED) [1, 2], which may be due to life-threatening myocardial infarction (MI) or benign musculoskeletal pain [3]. Majority of chest pain patients are subjected to extensive diagnostic tests to rule out acute coronary syndrome (ACS), resulting in oftentimes, unnecessary prolonged and costly ED admission, since only a small proportion of these patients will eventually receive a diagnosis of ACS [3]. Hence, early identification of chest pain patients at high-risk of developing adverse cardiac events has been a pressing issue to contend with in the ED. Several established clinical scores have been used for risk stratifying chest pain patients in the ED [4, 5], including the History, ECG, Age, Risk factors and Troponin (HEART) [6], the Thrombolysis in Myocardial Infarction (TIMI) [7], and the Global Registry of Acute Coronary Events (GRACE) [8] scores. Of the available clinical scores, the HEART score is the most accurate and widely used [5, 9–12]. Further research has focused on developing risk score-based clinical pathways for safe discharge of low-risk patients [1, 3, 13, 14].

In a recent review of clinical scores for ED patients with chest pain [5], heart rate variability (HRV) provided an alternative approach to build predictive models for accurate risk stratification [15–17]. HRV is a widely adopted tool for evaluating changes in cardiac autonomic regulation, and has been shown to be strongly associated with the autonomic nervous system (ANS) [18–20]. HRV analysis characterizes the beat-to-beat variation in an electrocardiogram (ECG) by utilizing time domain, frequency domain, and nonlinear analyses [19]. Reduced HRV has been found to be a significant predictor of adverse cardiac outcomes [21]. Given the complexity of quantifying HRV representation, several tools such as the PhysioNet Cardiovascular Signal Toolbox [22] and Kubios HRV [23] have been developed to standardize HRV analyses.

Based on the principle of parameter calculation on normal R-R intervals (RRIs; in this paper, RRIs are equivalent to normal-to-normal [NN] intervals, in which abnormal beats have been removed), HRV analysis generates only one set of parameters from a fixed length of ECG record. This limits the amount of information that can be extracted from raw ECG signals. In this paper, we proposed a novel representation of beat-to-beat variation, named as heart rate n-variability (HRnV) [24] to characterize RRIs from a different perspective. With the use of HRnV measures, multiple sets of parameters are calculated from the same ECG record, which significantly increases the amount of extracted information. Our study is the first clinical application of the HRnV representation, in which the value of this novel measure is evaluated in risk stratification of chest pain patients in the ED. We hypothesize that HRnV is closely related to conventional HRV while providing supplementary information that is associated with adverse cardiac events. Further, we will investigate the potential use of HRnV parameters to develop risk prediction tools.

## Methods

### Study Design and Setting

We conducted a retrospective analysis of data collected in our previous study on risk stratification of chest pain patients in the ED [9]. A convenience sample of patients was recruited at the ED of Singapore General Hospital between September 2010 and July 2015. This is a tertiary hospital with around-the-clock primary percutaneous coronary intervention capabilities and a median door-to-balloon time of 101 minutes [25]. At ED triage, patients are assigned a Patient Acuity Category Scale (PACS) score ranging from 1 to 4 to indicate the urgency for consultation; smaller number means higher acuity. In this study, patients >20 years old who presented to the ED with chief complaint of chest pain and were triaged to PACS 1 or 2 were included. Patients were excluded if they had ST-elevation myocardial infarction (STEMI) or an obvious non-cardiac etiology of chest pain diagnosed by the primary emergency physician. Patients were also excluded if their ECGs had high percentage of noise or they were in non-sinus rhythm; these criteria were applied to ensure the quality of HRV and HRnV analyses. The ethical approval was obtained from the Centralized Institutional Review Board (CIRB, Ref: 2014/584/C) of SingHealth, the largest public healthcare system in Singapore that includes the Singapore General Hospital as a key partner. In the ethical approval, patient consent was waived.

### Data Collection

During the data collection period, five to six-minute single-lead (lead II) ECG tracings were retrieved from the X-Series Monitor (ZOLL Medical Corporation, Chelmsford, MA). The first set of vital signs and troponin values from the recruited patients were extracted from the hospital’s electronic health records (EHR). In this study, high-sensitivity troponin-T was used, and an abnormal value was defined as >0.03 ng/mL [26]; it was further stratified into three groups and coded as 0 if the value was ≤0.03 ng/mL, 1 if the value was between 1 and 3 times the normal limit, and 2 if the value was >3 times the normal limit. Additionally, the patients’ first 12-lead ECGs were interpreted by two independent clinical reviewers. Pathologic ST-elevation, ST-depression, T-wave inversions, and Q-waves were recorded. Patient demographics, medical history, and information required for computing the HEART, TIMI, and GRACE scores were retrospectively reviewed and obtained from EHR.

### Proposed HRnV Representation of Beat-to-Beat Variation in ECG

#### HR_*n*_V: A Novel Measure with Non-Overlapped RRIs

Prior to introducing the new HR_*n*_V measure, we define a new type of RRI called RR*n*I, where 1 ≤ *n* ≤ *N*, and 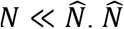 is the total number of RRIs. The definition of RR_*n*_I is illustrated in Figure 1a. When *n* = 1, RR_*n*_I becomes conventional RRI, that is, RR_1_I is equal to RRI. When *n* > 1, every *n* adjacent RRI is connected to form a new sequence of RR_*n*_Is. By using this strategy, we can create a maximum number of (*N* − 1) new RR_*n*_I sequences from conventional single RRI sequence. With these newly generated RR_*n*_I sequences, the calculation of HR_*n*_V parameters is straightforward and can be accomplished by applying established quantitative methods including time domain analysis, frequency domain analysis, and nonlinear analysis [18, 19]. In describing this new measure, we use the term “HR_*n*_V” prior to parameter names to indicate that these parameters are calculated from RR_*n*_I sequences. As noted in the above, HR_*n*_V is a novel measure based on newly generated, non-overlapped RR_*n*_Is. The computed HR_*n*_V parameters include but are not limited to the following: the average of RR_*n*_Is (HR_*n*_V mean NN), standard deviation of RR_*n*_Is (HR_*n*_V SDNN), square root of the mean squared differences between RR_*n*_Is (HR_*n*_V RMSSD), the number of times that the absolute difference between two successive RR_*n*_Is exceeds 50 ms (HR_*n*_V NN50), HR_*n*_V NN50 divided by the total number of RR_*n*_Is (HR_*n*_V pNN50), the integral of the RR_*n*_I histogram divided by the height of the histogram (HR_*n*_V triangular index), low frequency power (HR_*n*_V LF power), high frequency power (HR_*n*_V HF power), approximate entropy (HR_*n*_V ApEn), sample entropy (HR_*n*_V SampEn), and detrended fluctuation analysis (HR_*n*_V DFA), among others. Notably, two new parameters NN50*n* and pNN50*n* are created, where 50×*n* ms is set as the threshold to assess the difference between pairs of consecutive RR_*n*_Is.

**Figure 1:**
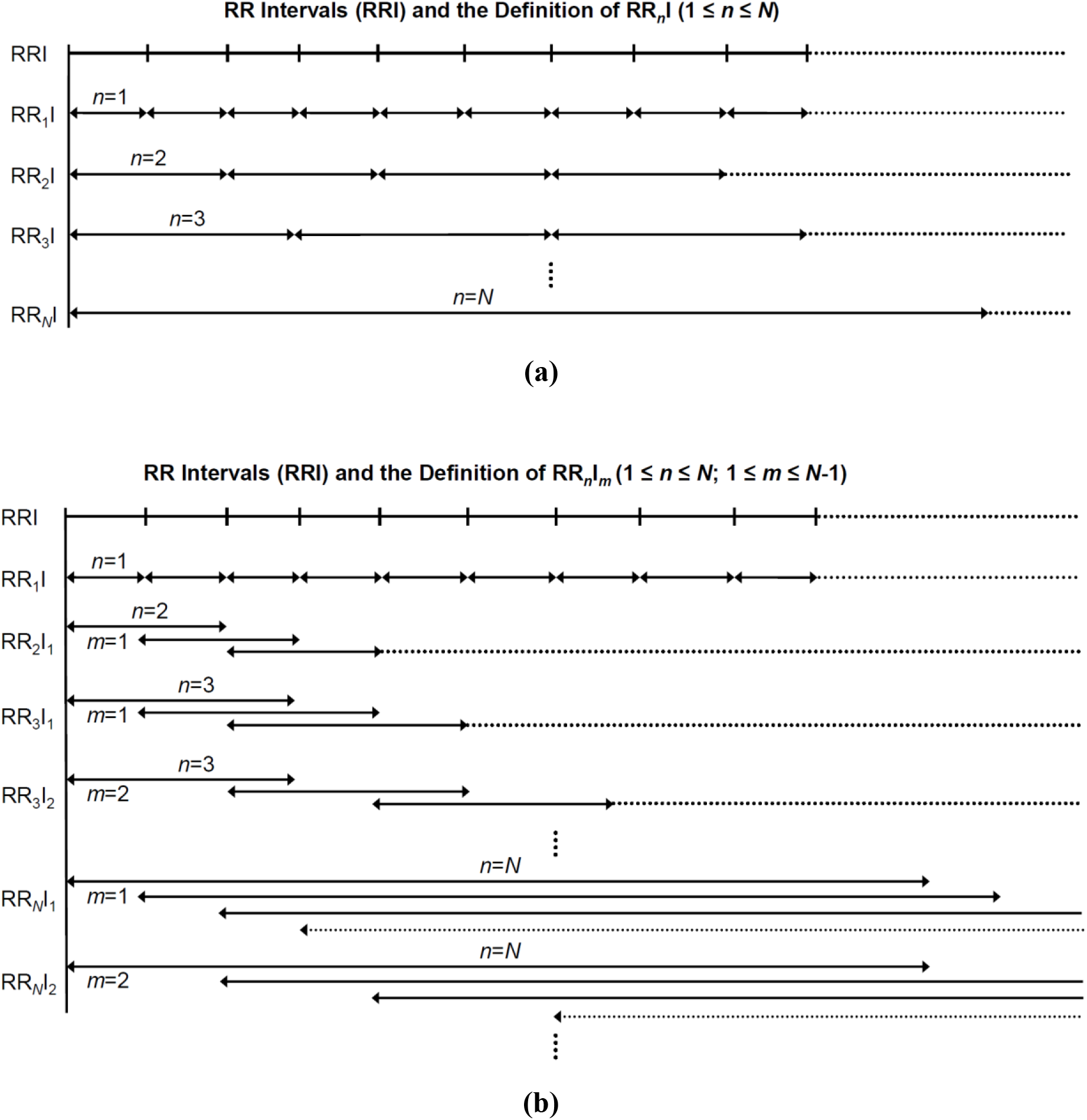
**(a)** Illustration of R-R intervals (RRIs) and the definition of RR_*n*_I where 1 ≤ *n* ≤ *N* and 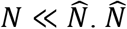 is the total number of RRIs; **(b)** Illustration of RRIs and the definition of RR_*n*_I_*m*_ where 1 ≤ *n* ≤ *N*, 1 ≤ *m* ≤ *N* − 1, and 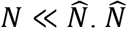 is the total number of RRIs and *m* indicates the non-overlapped portion between two consecutive RR_*n*_I_*m*_ sequences.

**Figure 2:**
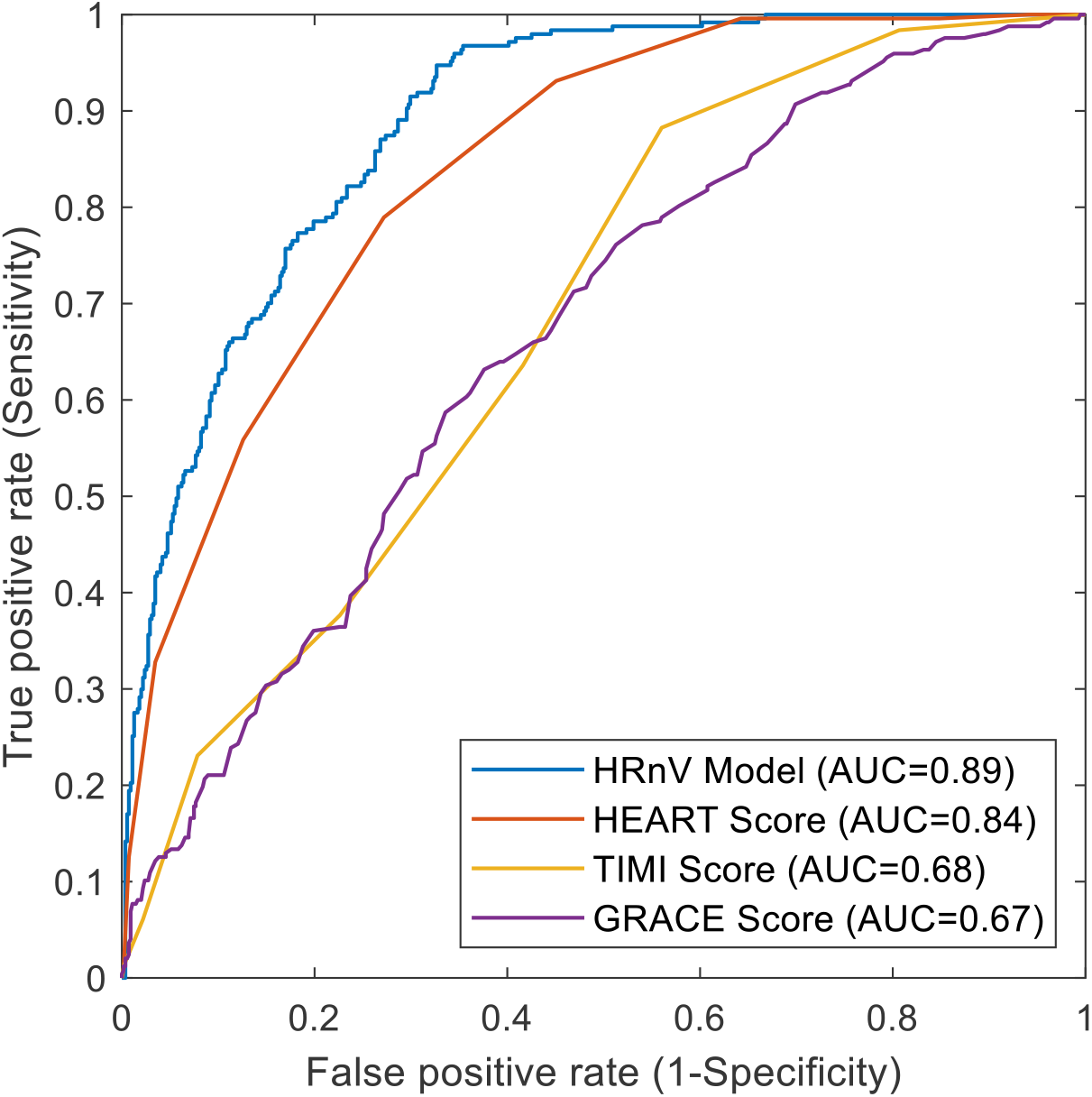
The receiver operating characteristic (ROC) curves produced by the heart rate n-variability (HRnV) model (performance was based on leave-one-out cross-validation), the History, ECG, Age, Risk factors and Troponin (HEART) score, the Thrombolysis in Myocardial Infarction (TIMI) score, and the Global Registry of Acute Coronary Events (GRACE) score.

#### HR_*n*_V_*m*_: A Novel Measure with Overlapped RRIs

Like RR_*n*_I that is used in HR_*n*_V, to define the HR_*n*_V_*m*_ measure we introduce another type of RRI called RR_*n*_I_*m*_, where 1 ≤ *n* ≤ *N*, 1 ≤ *m* ≤ *N* − 1, and 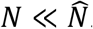. In the RR_*n*_I_*m*_ sequence, *m* is used to indicate the level of overlap between consecutive RR_*n*_I_*m*_ sequences. As illustrated in Figure 1b, (*n* − *m*) RRIs form the overlapped portions. When *m* = *n*, RR_*n*_I_*m*_ becomes RR_*n*_I; therefore, the upper limit of *m* is *N* − 1. By controlling the overlap among these newly generated RR_*n*_I_*m*_ sequences, we can create a maximum number of (*N* × (*N* − 1)/2) RR_*n*_I_*m*_ sequences (excluding the RR_*n*_I sequence) from conventional single RRI sequence. For each of the newly created RR_*n*_I_*m*_ sequences, we apply time domain analysis, frequency domain analysis, and nonlinear analysis to calculate HR_*n*_V_*m*_ parameters. We add the term “HR_*n*_V_*m*_” prior to the parameters to denote that they are computed from RR_*n*_I_*m*_ sequences. For example, the average RR_*n*_I_*m*_ intervals and the sample entropy are written as HR_*n*_V_*m*_ mean NN and HR_*n*_V_*m*_ SampEn, respectively. The HR_*n*_V_*m*_ measure extracts additional information than HR_*n*_V does, by adopting a strategy of controlling sequence overlap.

### HRnV Analysis and Parameter Calculation

We developed the HRnV-Calc software suite (https://github.com/HRnV) to calculate HRnV parameters. The HRnV-Calc software integrates functions from the PhysioNet Cardiovascular Signal Toolbox [22] to perform standardized ECG signal processing and QRS complex detection. Given the short ECG records in this study, the upper limit of *n* was set as three; thus, six sets of parameters were calculated, namely HRV, HR_2_V, HR_2_V_1_, HR_3_V, HR_3_V_1_, and HR_3_V_2_.

### Clinical Outcomes

The primary endpoint in this study was a composite outcome of major adverse cardiac events (MACE) [27], including all-cause death, acute myocardial infarction (AMI), and revascularization (coronary artery bypass graft [CABG] or percutaneous coronary intervention [PCI]) within 30 days of ED presentation.

### Statistical Analysis

Continuous variables were presented in terms of mean and standard deviation and compared between the two categories of the primary outcome (MACE) using two-sample t-test. Categorical variables were presented in terms of frequency and percentage and compared between the two categories of the primary outcome (MACE) using chi-square test. A statistically significant difference was defined as p<0.05. To evaluate the HRnV parameters and other risk factors, we conducted univariable and multivariable analyses and subsequently developed simple prediction models using traditional logistic regression. In building the HRnV prediction model, we selected candidate variables with p<0.2 in the univariable analysis and fed them into multivariable stepwise logistic regression. To evaluate the predictive performance, we used leave-one-out cross-validation (LOOCV) to conduct the analysis.

The receiver operating characteristic (ROC) analysis [28] was performed to compare prediction performances among the HRnV model, HEART, TIMI and GRACE scores. The area under the ROC curve (AUC), sensitivity, specificity, positive predictive value (PPV), and negative predictive value (NPV) were reported as the predictive measures. Data preparation, descriptive analysis, and predictive model development were performed in R version 3.6.0 (R Foundation, Vienna, Austria), and the ROC analysis was conducted in MATLAB R2019a (MathWorks, Natick, MA).

## Results

A total of 795 patients were selected from the originally recruited 922 patients [9]. 28 were excluded for ECG recording issues, four were excluded for clearly non-cardiac chest pain, and 95 were excluded for irregular rhythm/artifacts. Among the included 795 patients, 247 (31%) met the primary outcome of 30-day MACE. Table 1 shows patient baseline characteristics. Patients with the primary outcome were older (mean age 61 years vs. 59 years, p=0.035) and had a higher proportion of males (76.1% vs. 64.6%, p=0.002). There were no statistically significant differences between MACE and non-MACE groups in terms of patient ethnicity. Factors such as history of diabetes and current smoking status were significantly more prevalent in the group with MACE.

**Table 1:** Patient baseline characteristics.

|  | **Total (n=795)** | **MACE**<br>**(n=247)** | **Non-MACE**<br>**(n=548)** | **p-value** |
| --- | --- | --- | --- | --- |
| Age, mean (SD) | 59.63 (12.88) | 61.06 (11.38) | 58.99 (13.47) | 0.035 |
| Male gender, n (%) | 542 (68.2) | 188 (76.1) | 354 (64.6) | 0.002 |
| Race, n (%) |  |  |  | 0.623 |
| Chinese | 492 (61.9) | 159 (64.4) | 333 (60.8) |  |
| Indian | 129 (16.2) | 34 (13.8) | 95 (17.3) |  |
| Malay | 150 (18.9) | 46 (18.6) | 104 (19.0) |  |
| Other | 24 (3.0) | 8 (3.2) | 16 (2.9) |  |
| Medical history, n (%) |  |  |  |  |
| Ischemic heart disease | 343 (43.1) | 115 (46.6) | 228 (41.6) | 0.22 |
| Diabetes | 278 (35.0) | 106 (42.9) | 172 (31.4) | 0.002 |
| Hypertension | 509 (64.0) | 161 (65.2) | 348 (63.5) | 0.707 |
| Hypercholesterolemia | 476 (59.9) | 151 (61.1) | 325 (59.3) | 0.683 |
| Stroke | 58 (7.3) | 15 (6.1) | 43 (7.8) | 0.458 |
| Cancer | 29 (3.6) | 7 (2.8) | 22 (4.0) | 0.537 |
| Respiratory disease | 31 (3.9) | 5 (2.0) | 26 (4.7) | 0.102 |
| Chronic kidney disease | 87 (10.9) | 26 (10.5) | 61 (11.1) | 0.32 |
| Congestive heart failure | 38 (4.8) | 9 (3.6) | 29 (5.3) | 0.407 |
| History of PCI | 199 (25.0) | 68 (27.5) | 131 (23.9) | 0.316 |
| History of CABG | 71 (8.9) | 26 (10.5) | 45 (8.2) | 0.355 |
| History of AMI | 133 (16.7) | 48 (19.4) | 85 (15.5) | 0.288 |
| Active smoker | 197 (24.8) | 73 (29.6) | 124 (22.6) | 0.003 |

MACE, major adverse cardiac events; SD, standard deviation; PCI, percutaneous coronary intervention; CABG, coronary artery bypass graft; AMI, acute myocardial infarction.

Descriptive analyses of HRV and HRnV parameters are tabulated in Table 2. In this clinical case study, *N* was set as 3, thus HR_2_V, HR_2_V_1_, HR_3_V, HR_3_V_1_ and HR_3_V_2_ parameters were calculated. Among time domain parameters such as mean NN, SDNN and RMSSD, the HR_*n*_V and HR_*n*_V_*m*_ values were generally incremental with an increase in *n*. Notably, HR_2_V NN50 and HR_3_V NN50 were much lower than conventional HRV NN50. Moreover, NN50*n* and pNN50*n* are parameters specifically applicable to the HRnV representation. Like time domain parameters, the same trend of changes in frequency domain parameters were observed. The magnitude of increment in VLF power and LF power was larger than that of HF power with increasing *n*. However, one exception was the normalized HF power, whose HR_*n*_V and HR_*n*_V_*m*_ parameters were smaller than that of HRV. In nonlinear analysis, the differences in Poincaré SD2 values were obvious between HRV and HRnV parameters. HR_2_V SampEn and HR_3_V SampEn were remarkably larger compared to SampEn parameters of HRV, HR_2_V_1_, HR_3_V_1_, and HR_3_V_2_, as the confidence interval of SampEn was wide when data points were less than 200 [19], since our ECG recordings were only five to six-minute long. HR_2_V_1_, HR_3_V_1_ and HR_3_V_2_ were free from this issue as they were calculated from overlapping RR_*n*_I_*m*_ sequences where more than 200 data points were available.

**Table 2:** Descriptive analyses of heart rate variability (HRV) and heart rate n-variability (HRnV) parameters.

|  | HRV | HR<sub>2</sub>V | HR<sub>2</sub>V<sub>1</sub> | HR<sub>3</sub>V | HR<sub>3</sub>V<sub>1</sub> | HR<sub>3</sub>V<sub>2</sub> |
| --- | --- | --- | --- | --- | --- | --- |
| Mean NN (s) | 829.40 (169.49) | 1656.65 (339.85) | 1658.81 (338.99) | 2484.80 (509.33) | 2488.22 (508.50) | 2485.02 (509.84) |
| SDNN (s) | 38.16 (25.49) | 62.28 (45.45) | 68.81 (47.00) | 82.06 (62.47) | 97.79 (67.46) | 87.77 (64.52) |
| RMSSD (s) | 30.04 (23.07) | 32.61 (26.68) | 33.79 (25.67) | 34.83 (28.86) | 36.27 (26.50) | 34.98 (27.43) |
| Skewness | -0.65 (2.34) | -0.41 (1.66) | -0.59 (1.95) | -0.29 (1.29) | -0.55 (1.69) | -0.38 (1.42) |
| Kurtosis | 14.59 (26.83) | 7.33 (13.58) | 10.17 (17.90) | 5.15 (8.13) | 8.06 (12.92) | 5.98 (9.75) |
| Triangular index | 7.68 (4.19) | 10.38 (5.10) | 12.60 (6.45) | 11.47 (5.29) | 16.25 (7.94) | 13.06 (6.04) |
| NN50 (count) | 21.08 (33.98) | 14.46 (20.35) | 29.35 (40.03) | 11.57 (15.05) | 35.29 (44.34) | 17.41 (22.51) |
| pNN50 (%) | 6.31 (11.08) | 8.66 (13.18) | 8.75 (12.97) | 10.31 (14.27) | 10.38 (13.95) | 10.28 (14.20) |
| NN50<sub>n</sub> (count) | - | 4.16 (9.72) | 8.45 (18.76) | 1.37 (3.72) | 4.37 (10.72) | 2.08 (5.48) |
| pNN50<sub>n</sub> (%) | - | 2.60 (6.67) | 2.64 (6.47) | 1.32 (3.95) | 1.39 (3.86) | 1.33 (3.87) |
| Total power (ms<sup>2</sup>) | 2518.30 (4797.05) | 7797.46<br>(16947.44) | 9156.26<br>(17970.75) | 13904.78<br>(37182.24) | 18714.67<br>(37620.26) | 15706.11<br>(34845.52) |
| VLF power (ms<sup>2</sup>) | 985.18 (1991.52) | 3401.42 (6569.37) | 3922.74 (7987.46) | 6503.53<br>(14205.11) | 8772.26<br>(17986.63) | 7567.79<br>(14666.32) |
| LF power (ms<sup>2</sup>) | 732.36 (1841.88) | 2626.83 (7593.16) | 2782.48 (7212.62) | 5091.49<br>(18402.20) | 5740.99<br>(15243.38) | 5397.76<br>(16001.18) |
| HF power (ms<sup>2</sup>) | 527.27 (1232.69) | 1328.86 (4033.96) | 1361.53 (3433.55) | 1661.69 (7237.55) | 1762.45 (4851.11) | 1761.05 (6477.63) |
| LF power norm<br>(nu) | 56.76 (19.20) | 66.82 (18.17) | 66.42 (17.35) | 76.53 (15.32) | 77.65 (14.55) | 77.93 (14.95) |
| HF power norm<br>(nu) | 43.24 (19.20) | 33.18 (18.17) | 33.58 (17.35) | 23.47 (15.32) | 22.35 (14.55) | 22.07 (14.95) |
| LF/HF | 1.99 (1.93) | 3.24 (2.95) | 3.04 (2.73) | 5.60 (5.21) | 5.79 (4.99) | 6.06 (5.18) |
| Poincaré SD1<br>(ms) | 21.27 (16.34) | 23.12 (18.93) | 23.92 (18.18) | 24.72 (20.50) | 25.68 (18.77) | 24.80 (19.46) |
| Poincaré SD2<br>(ms) | 48.82 (33.29) | 84.47 (62.15) | 93.88 (64.58) | 112.87 (86.62) | 135.55 (94.02) | 121.20 (89.72) |
| SampEn | 1.57 (0.51) | 83.84 (2324.24) | 1.33 (0.48) | 248.48 (4020.64) | 1.06 (0.41) | 1.14 (0.45) |
| ApEn | 0.99 (0.20) | 0.72 (0.18) | 0.91 (0.17) | 0.60 (0.15) | 0.84 (0.17) | 0.70 (0.15) |
| DFA, $\alpha_1$ | 0.99 (0.31) | 1.24 (0.29) | 1.23 (0.27) | 1.41 (0.27) | 1.42 (0.23) | 1.42 (0.25) |
| DFA, $\alpha_2$ | 0.95 (0.22) | 0.98 (0.35) | 0.98 (0.22) | 0.86 (0.65) | 1.01 (0.22) | 1.02 (0.36) |

HRV, heart rate variability; mean NN, average of R-R intervals; SDNN, standard deviation of R-R intervals; RMSSD, square root of the mean squared differences between R-R intervals; NN50, the number of times that the absolute difference between 2 successive R-R intervals exceeds 50 ms; pNN50, NN50 divided by the total number of R-R intervals; NN50<sub>n</sub>, the number of times that the absolute difference between 2 successive RR<sub>n</sub>I/RR<sub>n</sub>I<sub>m</sub> sequences exceeds 50×*n* ms; pNN50<sub>n</sub>, NN50<sub>n</sub> divided by the total number of RR<sub>n</sub>I/RR<sub>n</sub>I<sub>m</sub> sequences; VLF, very low frequency; LF, low frequency; HF, high frequency; SD, standard deviation; SampEn, sample entropy; ApEn, approximate entropy; DFA, detrended fluctuation analysis.

Table 3 presents the results of univariable analyses of HR_*n*_V and HR_*n*_V_*m*_ parameters. Eleven out of 21 conventional HRV parameters were statistically significant. Additionally, 13 HR_2_V, six HR_3_V, 11 HR_2_V_1_, seven HR_3_V_1_ and 11 HR_3_V_2_ parameters were also significant. Overall, additional 115 HRnV parameters were derived, among which 48 showed statistical significance between patients who had 30-day MACE and who did not. Among all HRV and HRnV parameters, mean NN, SDNN, RMSSD, NN50, pNN50, HF power, Poincaré SD1 and SD2 appeared statistically significant in at least five out of six measures (i.e., HRV, HR_2_V, HR_2_V_1_, HR_3_V, HR_3_V_1_, and HR_3_V_2_). Furthermore, skewness, LF power, SampEn, and ApEn, which did not demonstrate statistical significance in conventional HRV analysis, became statistically significant in HRnV representation. Table 4 presents the results of multivariable analyses on HR_*n*_V and HR_*n*_V_*m*_ parameters by adjusting for age and sex. Compared with the results in Table 3, several parameters became statistically insignificant, e.g. NN50 of HR_3_V and HR_3_V_2_, and triangular index of HRV, HR_2_V, and HR_3_V_2_. On the other hand, ApEn of HR_2_V, HR_2_V_1_, and HR_3_V_2_ became statistically significant after the adjustment.

**Table 3:** Univariable analysis of HR_*n*_V and HR_*n*_V_*m*_ parameters.

|  | **HRV** |  | **HR<sub>2</sub>V** |  | **HR<sub>3</sub>V** |  |
| --- | --- | --- | --- | --- | --- | --- |
|  | OR (95% CI) | p | OR (95% CI) | p | OR (95% CI) | p |
| Mean NN | 0.999 (0.998-1.000) | 0.023* | 0.999 (0.999-1.000) | 0.023* | 1.000 (0.999-1.000) | 0.023* |
| SDNN | 0.992 (0.986-0.999) | 0.023* | 0.996 (0.992-1.000) | 0.028* | 0.997 (0.995-1.000) | 0.060 |
| RMSSD | 0.990 (0.982-0.998) | 0.010* | 0.992 (0.985-0.998) | 0.011* | 0.994 (0.988-0.999) | 0.030* |
| Skewness | 1.059 (0.991-1.132) | 0.088 | 1.079 (0.981-1.186) | 0.118 | 1.139 (1.006-1.290) | 0.040* |
| Kurtosis | 1.006 (1.000-1.011) | 0.038* | 1.009 (0.998-1.019) | 0.113 | 1.011 (0.993-1.029) | 0.242 |
| Triangular index | 0.961 (0.925-0.998) | 0.039* | 0.967 (0.938-0.997) | 0.032* | 0.978 (0.950-1.007) | 0.133 |
| NN50 | 0.993 (0.987-0.998) | 0.008* | 0.989 (0.981-0.998) | 0.012* | 0.988 (0.977-0.999) | 0.031* |
| pNN50 | 0.978 (0.962-0.995) | 0.009* | 0.984 (0.971-0.997) | 0.014* | 0.987 (0.976-0.999) | 0.027* |
| NN50<sub>n</sub> | - | - | 0.982 (0.964-1.001) | 0.065 | 0.952 (0.905-1.002) | 0.059 |
| pNN50<sub>n</sub> | - | - | 0.974 (0.946-1.002) | 0.069 | 0.951 (0.903-1.001) | 0.054 |
| Total power | 1.000 (1.000-1.000) | 0.031* | 1.000 (1.000-1.000) | 0.021* | 1.000 (1.000-1.000) | 0.072 |
| VLF power | 1.000 (1.000-1.000) | 0.132 | 1.000 (1.000-1.000) | 0.070 | 1.000 (1.000-1.000) | 0.133 |
| LF power | 1.000 (1.000-1.000) | 0.077 | 1.000 (1.000-1.000) | 0.023* | 1.000 (1.000-1.000) | 0.063 |
| HF power | 1.000 (0.999-1.000) | 0.002* | 1.000 (1.000-1.000) | 0.014* | 1.000 (1.000-1.000) | 0.074 |
| LF power norm | 1.001 (0.994-1.009) | 0.738 | 0.999 (0.99-1.007) | 0.733 | 0.994 (0.985-1.004) | 0.248 |
| HF power norm | 0.999 (0.991-1.007) | 0.738 | 1.001 (0.993-1.01) | 0.733 | 1.006 (0.996-1.015) | 0.248 |
| LF/HF | 1.034 (0.959-1.116) | 0.381 | 1.014 (0.964-1.066) | 0.592 | 1.001 (0.973-1.031) | 0.923 |
| Poincaré SD1 | 0.986 (0.975-0.997) | 0.010* | 0.988 (0.979-0.997) | 0.011* | 0.991 (0.983-0.999) | 0.029* |
| Poincaré SD2 | 0.995 (0.990-1.000) | 0.032* | 0.997 (0.994-1.000) | 0.032* | 0.998 (0.996-1.000) | 0.063 |
| SampEn | 0.813 (0.604-1.095) | 0.173 | 0.730 (0.545-0.977) | 0.035* | 1.000 (1.000-1.000) | 0.932 |
| ApEn | 1.645 (0.752-3.598) | 0.213 | 2.319 (1.003-5.357) | 0.049* | 1.241 (0.463-3.327) | 0.667 |
| DFA, $\alpha_1$ | 0.953 (0.585-1.552) | 0.846 | 1.031 (0.611-1.741) | 0.908 | 0.968 (0.560-1.672) | 0.907 |
| DFA, $\alpha_2$ | 1.532 (0.773-3.034) | 0.221 | 1.202 (0.782-1.848) | 0.401 | 1.184 (0.934-1.500) | 0.163 |
|  | **HR<sub>2</sub>V<sub>1</sub>** |  | **HR<sub>3</sub>V<sub>1</sub>** |  | **HR<sub>3</sub>V<sub>2</sub>** |  |
|  | OR (95% CI) | p | OR (95% CI) | p | OR (95% CI) | p |
| Mean NN | 0.999 (0.999-1.000) | 0.023* | 1.000 (0.999-1.000) | 0.023* | 1.000 (0.999-1.000) | 0.023* |
| SDNN | 0.996 (0.993-1.000) | 0.034* | 0.997 (0.995-1.000) | 0.042* | 0.997 (0.995-1.000) | 0.034* |
| RMSSD | 0.991 (0.984-0.998) | 0.010* | 0.992 (0.986-0.999) | 0.016* | 0.993 (0.986-0.999) | 0.016* |
| Skewness | 1.061 (0.980-1.149) | 0.144 | 1.072 (0.978-1.176) | 0.139 | 1.098 (0.982-1.227) | 0.100 |
| Kurtosis | 1.007 (0.999-1.015) | 0.082 | 1.006 (0.994-1.017) | 0.333 | 1.010 (0.995-1.025) | 0.195 |
| Triangular index | 0.981 (0.958-1.005) | 0.119 | 0.982 (0.963-1.001) | 0.065 | 0.974 (0.949-0.999) | 0.040* |
| NN50 | 0.995 (0.991-0.999) | 0.018* | 0.996 (0.993-1.000) | 0.052 | 0.992 (0.985-0.999) | 0.035* |
| pNN50 | 0.984 (0.972-0.997) | 0.020* | 0.988 (0.977-1.000) | 0.049* | 0.988 (0.976-0.999) | 0.035* |
| NN50<sub>n</sub> | 0.989 (0.979-1.000) | 0.043* | 0.982 (0.964-1.000) | 0.054 | 0.974 (0.943-1.007) | 0.118 |
| pNN50<sub>n</sub> | 0.969 (0.939-0.999) | 0.046* | 0.947 (0.895-1.002) | 0.058 | 0.960 (0.914-1.009) | 0.109 |
| Total power | 1.000 (1.000-1.000) | 0.048* | 1.000 (1.000-1.000) | 0.072 | 1.000 (1.000-1.000) | 0.029* |
| VLF power | 1.000 (1.000-1.000) | 0.139 | 1.000 (1.000-1.000) | 0.145 | 1.000 (1.000-1.000) | 0.074 |
| LF power | 1.000 (1.000-1.000) | 0.084 | 1.000 (1.000-1.000) | 0.092 | 1.000 (1.000-1.000) | 0.027* |
| HF power | 1.000 (1.000-1.000) | 0.005* | 1.000 (1.000-1.000) | 0.010* | 1.000 (1.000-1.000) | 0.022* |
| LF power norm | 1.000 (0.991-1.008) | 0.937 | 0.995 (0.985-1.006) | 0.382 | 0.995 (0.986-1.005) | 0.356 |
| HF power norm | 1.000 (0.992-1.009) | 0.937 | 1.005 (0.994-1.015) | 0.382 | 1.005 (0.995-1.015) | 0.356 |
| LF/HF | 1.024 (0.970-1.080) | 0.387 | 1.003 (0.973-1.033) | 0.863 | 0.999 (0.971-1.029) | 0.966 |
| Poincaré SD1 | 0.987 (0.978-0.997) | 0.010* | 0.989 (0.980-0.998) | 0.016* | 0.989 (0.981-0.998) | 0.016* |
| Poincaré SD2 | 0.997 (0.995-1.000) | 0.039* | 0.998 (0.996-1.000) | 0.045* | 0.998 (0.996-1.000) | 0.037* |
| SampEn | 0.854 (0.623-1.171) | 0.328 | 0.802 (0.553-1.161) | 0.242 | 0.709 (0.500-1.005) | 0.053 |
| ApEn | 2.065 (0.842-5.064) | 0.113 | 1.207 (0.499-2.922) | 0.677 | 2.558 (0.906-7.222) | 0.076 |
| DFA, $\alpha_1$ | 0.888 (0.514-1.537) | 0.672 | 1.039 (0.547-1.971) | 0.907 | 1.004 (0.549-1.835) | 0.991 |
| DFA, $\alpha_2$ | 1.557 (0.782-3.098) | 0.208 | 1.554 (0.780-3.093) | 0.210 | 1.169 (0.764-1.789) | 0.472 |

HRV, heart rate variability; OR, odds ratio; CI, confidence interval; mean NN, average of R-R intervals; SDNN, standard deviation of R-R intervals; RMSSD, square root of the mean squared differences between R-R intervals; NN50, the number of times that the absolute difference between 2 successive R-R intervals exceeds 50 ms; pNN50, NN50 divided by the total number of R-R intervals; NN50 $n$ , the number of times that the absolute difference between 2 successive RR $_n$ I/RR $_n$ I $_m$ sequences exceeds 50 $\times n$ ms; pNN50 $n$ , NN50 $n$ divided by the total number of RR $_n$ I/RR $_n$ I $_m$ sequences; VLF, very low frequency; LF, low frequency; HF, high frequency; SD, standard deviation; SampEn, sample entropy; ApEn, approximate entropy; DFA, detrended fluctuation analysis.

**Table 4:** Multivariable analysis of HR_*n*_V and HR_*n*_V_*m*_ parameters by adjusting for age and sex.

|  | **HRV** |  | **HR<sub>2</sub>V** |  | **HR<sub>3</sub>V** |  |
| --- | --- | --- | --- | --- | --- | --- |
|  | OR (95% CI) | p | OR (95% CI) | p | OR (95% CI) | p |
| Mean NN | 0.999 (0.998-1) | 0.005* | 0.999 (0.999-1.000) | 0.005* | 1.000 (0.999-1.000) | 0.005* |
| SDNN | 0.993 (0.986-0.999) | 0.035* | 0.996 (0.992-1.000) | 0.040* | 0.998 (0.995-1.000) | 0.093 |
| RMSSD | 0.990 (0.982-0.998) | 0.011* | 0.992 (0.985-0.999) | 0.016* | 0.994 (0.988-1.000) | 0.047* |
| Skewness | 1.064 (0.995-1.138) | 0.068 | 1.082 (0.983-1.191) | 0.109 | 1.140 (1.005-1.293) | 0.042* |
| Kurtosis | 1.005 (1.000-1.011) | 0.047* | 1.008 (0.997-1.019) | 0.139 | 1.011 (0.993-1.030) | 0.238 |
| Triangular index | 0.967 (0.93-1.006) | 0.093 | 0.971 (0.940-1.002) | 0.070 | 0.982 (0.953-1.013) | 0.256 |
| NN50 | 0.993 (0.988-0.999) | 0.013* | 0.991 (0.982-0.999) | 0.030* | 0.990 (0.979-1.001) | 0.078 |
| pNN50 | 0.979 (0.963-0.996) | 0.015* | 0.986 (0.972-0.999) | 0.033* | 0.989 (0.977-1.001) | 0.063 |
| NN50<sub>n</sub> | - | - | 0.983 (0.964-1.002) | 0.081 | 0.954 (0.906-1.005) | 0.077 |
| pNN50<sub>n</sub> | - | - | 0.975 (0.947-1.004) | 0.086 | 0.952 (0.903-1.004) | 0.069 |
| Total power | 1.000 (1.000-1.000) | 0.042* | 1.000 (1.000-1.000) | 0.026* | 1.000 (1.000-1.000) | 0.104 |
| VLF power | 1.000 (1.000-1.000) | 0.167 | 1.000 (1.000-1.000) | 0.082 | 1.000 (1.000-1.000) | 0.152 |
| LF power | 1.000 (1.000-1.000) | 0.093 | 1.000 (1.000-1.000) | 0.033* | 1.000 (1.000-1.000) | 0.105 |
| HF power | 1.000 (0.999-1.000) | 0.003* | 1.000 (1.000-1.000) | 0.016* | 1.000 (1.000-1.000) | 0.101 |
| LF power norm | 1.002 (0.994-1.011) | 0.589 | 0.999 (0.990-1.007) | 0.769 | 0.994 (0.984-1.003) | 0.202 |
| HF power norm | 0.998 (0.989-1.006) | 0.589 | 1.001 (0.993-1.010) | 0.769 | 1.006 (0.997-1.016) | 0.202 |
| LF/HF | 1.039 (0.961-1.124) | 0.336 | 1.013 (0.962-1.066) | 0.620 | 0.999 (0.970-1.028) | 0.928 |
| Poincaré SD1 | 0.986 (0.975-0.997) | 0.011* | 0.989 (0.980-0.998) | 0.016* | 0.992 (0.983-1.000) | 0.047* |
| Poincaré SD2 | 0.995 (0.990-1.000) | 0.050* | 0.997 (0.994-1.000) | 0.046* | 0.998 (0.996-1.000) | 0.098 |
| SampEn | 0.852 (0.630-1.152) | 0.297 | 0.752 (0.559-1.010) | 0.058 | 1.000 (1.000-1.000) | 0.956 |
| ApEn | 1.669 (0.754-3.693) | 0.207 | 2.668 (1.139-6.246) | 0.024* | 1.507 (0.555-4.096) | 0.421 |
| DFA, $\alpha_1$ | 0.991 (0.593-1.654) | 0.971 | 1.072 (0.622-1.848) | 0.802 | 0.962 (0.550-1.682) | 0.891 |
| DFA, $\alpha_2$ | 1.499 (0.750-2.993) | 0.252 | 1.204 (0.782-1.853) | 0.400 | 1.193 (0.941-1.512) | 0.146 |
|  | **HR<sub>2</sub>V<sub>1</sub>** |  | **HR<sub>3</sub>V<sub>1</sub>** |  | **HR<sub>3</sub>V<sub>2</sub>** |  |
|  | OR (95% CI) | p | OR (95% CI) | p | OR (95% CI) | p |
| Mean NN | 0.999 (0.999-1.000) | 0.005* | 1.000 (0.999-1.000) | 0.005* | 1.000 (0.999-1.000) | 0.005* |
| SDNN | 0.996 (0.993-1.000) | 0.052 | 0.998 (0.995-1.000) | 0.064 | 0.997 (0.995-1.000) | 0.049* |
| RMSSD | 0.992 (0.985-0.998) | 0.015* | 0.993 (0.986-0.999) | 0.023* | 0.993 (0.987-0.999) | 0.023* |
| Skewness | 1.066 (0.984-1.156) | 0.118 | 1.079 (0.983-1.185) | 0.108 | 1.099 (0.982-1.229) | 0.099 |
| Kurtosis | 1.007 (0.999-1.015) | 0.096 | 1.005 (0.994-1.017) | 0.377 | 1.010 (0.994-1.025) | 0.218 |
| Triangular index | 0.985 (0.960-1.010) | 0.234 | 0.985 (0.965-1.005) | 0.137 | 0.977 (0.951-1.003) | 0.088 |
| NN50 | 0.996 (0.991-1.000) | 0.047* | 0.997 (0.993-1.001) | 0.130 | 0.993 (0.986-1.001) | 0.084 |
| pNN50 | 0.986 (0.973-1.000) | 0.046* | 0.990 (0.979-1.002) | 0.111 | 0.989 (0.978-1.001) | 0.076 |
| NN50<sub>n</sub> | 0.990 (0.980-1.000) | 0.059* | 0.982 (0.963-1.001) | 0.064 | 0.975 (0.943-1.008) | 0.142 |
| pNN50<sub>n</sub> | 0.971 (0.941-1.002) | 0.063 | 0.947 (0.893-1.004) | 0.067 | 0.962 (0.915-1.012) | 0.131 |
| Total power | 1.000 (1.000-1.000) | 0.064 | 1.000 (1.000-1.000) | 0.096 | 1.000 (1.000-1.000) | 0.035* |
| VLF power | 1.000 (1.000-1.000) | 0.173 | 1.000 (1.000-1.000) | 0.180 | 1.000 (1.000-1.000) | 0.086 |
| LF power | 1.000 (1.000-1.000) | 0.100 | 1.000 (1.000-1.000) | 0.108 | 1.000 (1.000-1.000) | 0.037* |
| HF power | 1.000 (1.000-1.000) | 0.006* | 1.000 (1.000-1.000) | 0.014* | 1.000 (1.000-1.000) | 0.025* |
| LF power norm | 1.000 (0.991-1.009) | 0.960 | 0.995 (0.984-1.005) | 0.324 | 0.995 (0.985-1.005) | 0.329 |
| HF power norm | 1.000 (0.991-1.009) | 0.960 | 1.005 (0.995-1.016) | 0.324 | 1.005 (0.995-1.015) | 0.329 |
| LF/HF | 1.023 (0.968-1.081) | 0.428 | 0.999 (0.969-1.030) | 0.940 | 0.996 (0.967-1.026) | 0.786 |
| Poincaré SD1 | 0.988 (0.979-0.998) | 0.015* | 0.990 (0.981-0.999) | 0.023* | 0.990 (0.981-0.999) | 0.023* |
| Poincaré SD2 | 0.997 (0.995-1.000) | 0.059 | 0.998 (0.997-1.000) | 0.068 | 0.998 (0.996-1.000) | 0.052 |
| SampEn | 0.870 (0.632-1.197) | 0.393 | 0.842 (0.578-1.227) | 0.371 | 0.716 (0.504-1.019) | 0.064 |
| ApEn | 2.520 (1.009-6.298) | 0.048* | 1.413 (0.575-3.471) | 0.451 | 3.461 (1.201-9.971) | 0.021* |
| DFA, $\alpha_1$ | 0.898 (0.508-1.587) | 0.710 | 1.068 (0.555-2.058) | 0.843 | 1.005 (0.543-1.838) | 0.988 |
| DFA, $\alpha_2$ | 1.507 (0.751-3.025) | 0.249 | 1.500 (0.746-3.014) | 0.255 | 1.172 (0.764-1.798) | 0.467 |

HRV, heart rate variability; OR, odds ratio; CI, confidence interval; mean NN, average of R-R intervals; SDNN, standard deviation of R-R intervals; RMSSD, square root of the mean squared differences between R-R intervals; NN50, the number of times that the absolute difference between 2 successive R-R intervals exceeds 50 ms; pNN50, NN50 divided by the total number of R-R intervals; NN50 $n$ , the number of times that the absolute difference between 2 successive $RR_nI/RR_nI_m$ sequences exceeds 50 $\times n$ ms; pNN50 $n$ , NN50 $n$ divided by the total number of $RR_nI/RR_nI_m$ sequences; VLF, very low frequency; LF, low frequency; HF, high frequency; SD: standard deviation; SampEn, sample entropy; ApEn, approximate entropy; DFA: detrended fluctuation analysis.

Table 5 lists the 16 variables that were selected through multivariable stepwise logistic regression, among which there were one conventional HRV parameter and seven HRnV parameters. In addition to traditional predictors of adverse cardiac outcomes such as ST segment changes and troponin, HR_2_V ApEn (OR=0.095; 95% CI 0.014-0.628), HR_2_V_1_ ApEn (OR=19.700; 95% CI 2.942-131.900) and HR_3_V skewness (1.560; 95% CI 1.116-2.181) also demonstrated strong predictive power in assessing the risk of 30-day MACE. Table 6 presents the results of ROC analysis in evaluating the predictive performance by the HRnV model (based on LOOCV), HEART, TIMI, and GRACE scores. The HRnV model achieved the highest AUC value and outperformed HEART, TIMI, and GRACE scores in terms of specificity, PPV, and NPV at the optimal cut-off scores, defined as the points nearest to the upper-left corner of the ROC curves.

**Table 5:** Multivariable analysis with logistic regression on all variables.

| Variable | Adjusted OR | 95% CI |
| --- | --- | --- |
| Age | 1.021 | 1.002-1.041 |
| Diastolic BP | 1.018 | 1.003-1.034 |
| Pain score | 1.082 | 1.003-1.168 |
| ST-elevation | 6.449 | 2.762-15.059 |
| ST-depression | 4.827 | 2.511-9.277 |
| Q wave | 3.383 | 1.668-6.860 |
| Cardiac history† | 7.838 | 5.192-11.832 |
| Troponin | 4.406 | 3.218-6.033 |
| HRV NN50 | 0.981 | 0.970-0.991 |
| HR<sub>2</sub>V skewness | 0.806 | 0.622-1.045 |
| HR<sub>2</sub>V SampEn | 0.600 | 0.348-1.035 |
| HR<sub>2</sub>V ApEn | 0.095 | 0.014-0.628 |
| HR<sub>2</sub>V<sub>1</sub> ApEn | 19.700 | 2.942-131.900 |
| HR<sub>3</sub>V RMSSD | 1.024 | 1.008-1.040 |
| HR<sub>3</sub>V skewness | 1.560 | 1.116-2.181 |
| HR<sub>3</sub>V<sub>2</sub> HF power | 1.000 | 1.000-1.000 |

BP, blood pressure; HRV, heart rate variability; OR, odds ratio; CI, confidence interval; mean NN, average of R-R intervals; RMSSD, square root of the mean squared differences between R-R intervals; NN50, the number of times that the absolute difference between 2 successive R-R intervals exceeds 50 ms; LF, low frequency; HF, high frequency; SampEn, sample entropy; ApEn, approximate entropy.
† Cardiac history was a numeric value that was derived from the narrative in the hospital charts. Its value was zero if the patient history contained characteristics of atypical cardiac chest pain; Its value was two if the history contained characteristics of typical cardiac chest pain; Its value was one if the history contained characteristics of both atypical and typical cardiac chest pain.

**Table 6:** Comparison of performance of the HRnV model (based on leave-one-out cross-validation), HEART, TIMI, and GRACE scores in predicting 30-day major adverse cardiac events (MACE).

|  | AUC (95% CI) | Cut-off | Sensitivity (95% CI) | Specificity (95% CI) | PPV (95% CI) | NPV (95% CI) |
| --- | --- | --- | --- | --- | --- | --- |
| **HRnV Model** | 0.888 (0.860-0.917) | 0.3611† | 77.3% (72.1% - 82.5%) | 81.8% (78.5% - 85.0%) | 65.6% (60.2% - 71.1%) | 88.9% (86.1% - 91.6%) |
|  | - | 0.0352 | 99.2% (98.1% - 100.0%) | 39.6% (35.5% - 43.7%) | 42.5% (38.5% - 46.6%) | 99.1% (97.8% - 100.0%) |
| **HEART** | 0.841 (0.808-0.874) | 5† | 78.9% (73.9% - 84.0%) | 72.8% (69.1% - 76.5%) | 56.7% (51.4% - 61.9%) | 88.5% (85.5% - 91.4%) |
|  | - | 3 | 99.6% (98.8% - 100.0%) | 35.8% (31.8% - 39.8%) | 41.1% (37.2% - 45.1%) | 99.5% (98.5% - 100.0%) |
| **TIMI** | 0.681 (0.639-0.723) | 2† | 63.6% (57.6% - 69.6%) | 58.4% (54.3% - 62.5%) | 40.8% (35.9% - 45.7%) | 78.0% (74.0% - 82.1%) |
|  | - | 0 | 98.4% (96.8% - 100.0%) | 19.3% (16.0% - 22.7%) | 35.5% (31.9% - 39.1%) | 96.4% (92.9% - 99.9%) |
| **GRACE** | 0.665 (0.623-0.707) | 107† | 64.0% (58.0% - 70.0%) | 60.8% (56.7% - 64.9%) | 42.4% (37.3% - 47.4%) | 78.9% (75.0% - 82.8%) |
|  | - | 60 | 98.8% (97.4% - 100.0%) | 8.0% (5.8% - 10.3%) | 32.6% (29.3% - 36.0%) | 93.6% (86.6% - 100.0%) |

AUC, area under the curve; CI, confidence interval; PPV, positive predictive value; NPV, negative predictive value; HEART, History, ECG, Age, Risk factors and Troponin; TIMI, Thrombolysis in Myocardial Infarction; GRACE, Global Registry of Acute Coronary Events.
† Optimal cut-off values, defined as the points nearest to the upper-left corner on the ROC curves.

## Discussion

HRV has been well investigated in the past decades [18, 19, 29]. While the majority of efforts in HRV research were focused on development of advanced nonlinear techniques to derive novel parameters [30, 31], few investigated alternative approaches to analyze RRIs. Vollmer [32] used relative RRIs to describe the relative variation of consecutive RRIs, with which HRV was analyzed. Likewise, we proposed a novel HRnV representation, providing additional HRnV parameters than conventional HRV analysis. In this paper, we introduced two measures of HRnV, namely HR_*n*_V and HR_*n*_V_*m*_. HR_*n*_V is calculated based on non-overlapped RR_*n*_I sequences, while HR_*n*_V_*m*_ is computed from overlapped RR_*n*_I_*m*_ sequences. HRnV is not developed to replace the conventional HRV; instead, this representation is a natural extension of HRV. It enables the creation of additional parameters from raw ECGs, and thus empowers the extraction of supplementary information.

In our clinical case study, we investigated the predictive value of HRnV parameters to assess the risk of 30-day MACE for chest pain patients in the ED. In addition to 21 HRV parameters, 115 HRnV parameters were derived, of which 48 were found to be statistically significant in their association with the primary outcome. Notably, even with a small *n* (three in our study), newly generated HRnV parameters have greatly boosted the number of candidate predictors. When longer ECG records are available, more HRnV parameters can be calculated. With HRnV parameters, HRV parameters, vital signs, and several established risk factors, we conducted multivariable logistic regression analysis and selected age, diastolic BP, pain score, ST-elevation, ST-depression, Q wave, cardiac history, troponin, HRV NN50, and seven HRnV parameters. In addition to traditional risk factors like ST segment changes, HR_2_V ApEn, HR_2_V_1_ ApEn, and HR_3_V skewness were found as strong predictors for 30-day MACE. Comparing with the HEART, TIMI, and GRACE scores, the HRnV model achieved the highest AUC, specificity, PPV, and NPV values at the optimal cut-off points in ROC analysis. Therefore, HRnV may be clinically useful in determining the risk of 30-day MACE for ED patients with chest pain.

Due to the wide differential diagnosis for chest pain, accurate stratification is vital, particularly for preventing low-risk patients from obtaining expensive and unnecessary medical testing and intervention [3]. The TIMI and GRACE scores have been validated for risk prediction of patients with chest pain in the ED [4, 33, 34]. However, because they were originally developed for post-acute myocardial infarction, certain criteria within the TIMI and GRACE scores may be inappropriate for the undifferentiated chest pain cohorts in the ED [1]. In comparison, the HEART score was derived from a population of ED patients with chest pain, and has been extensively validated worldwide [10, 13, 27, 35]. It has proven its applicability in identifying both low-risk patients for possible early discharge and high-risk patients for urgent intervention. Built upon established scores, many chest pain pathways [14, 36–38] have been implemented and tested, particularly for the management of low-risk patients. Than et al. [38] evaluated a TIMI score-based accelerated diagnostic protocol (ADP) with a reported sensitivity of 99.3% and NPV of 99.1%. Similarly, a systematic review by Laureano-Phillips et al. [39] reported that the HEART score achieved both sensitivity and NPV of 100% in several validation studies. Furthermore, a cost-effectiveness study conducted in Brisbane, Australia reported economic benefits by adopting an ADP in the ED, with its associated reduction in expected cost and length of stay amongst patients with chest pain [40].

Most established clinical scores use conventional risk factors such as biomarkers, medical history, and presenting vital signs. However, patient history can sometimes be subjective and blood tests, such as troponin, take time to result. HRV, as a noninvasive measure, can be easily calculated from ECGs; it is an objective tool to assess the activities of the ANS [19]. It also has the advantage of requiring only several minutes to acquire (five to six minutes in our protocol), which is much faster than serum biomarkers. Over the past decades, HRV has been widely investigated in a broad range of clinical applications, particularly in cardiovascular research. HRV was found to be closely associated with sudden cardiac death [18]. It also showed significant correlations with adverse clinical outcomes in prehospital setting [41] and with MACE outcomes in ED patients with chest pain [17]. HRV parameters have been integrated with other risk factors into machine learning algorithms to predict adverse outcomes [42, 43]. These promising results motivated the use of HRV to develop objective and computerized risk stratification tools for chest pain patients [44, 45]. In an updated review of clinical scores for chest pain, Liu et al. [5] summarized several studies which aimed to develop alternative techniques for risk stratification.

This study aimed to present the novel HRnV representation and its measures and investigate their association with clinical outcomes. Although the HRnV parameters showed promising performance in identifying high-risk chest pain patients, this study was not intended to create a ready-to-use clinical tool. Instead, we have demonstrated the feasibility of utilizing HRnV parameters to augment conventional HRV and risk factors in designing a prediction tool/score. These HRnV parameters can be readily calculated without the collection of supplementary data. In this study, with five to six-minute ECG and *n* = 3, five-fold more HRnV parameters were calculated compared to HRV alone. When longer ECGs are available and parameter *n* is set as a larger number, more HRnV parameters can be derived. To build a HRnV-based risk stratification tool, a systematic approach is needed to derive a point-based, consistent score to ease its clinical application and practical implementation.

As a natural extension of conventional HRV, HRnV representation creates the opportunity to generate additional parameters. This representation could also serve as a smoother for RRIs, making them less sensitive to sudden changes caused by abnormal heart beats (e.g., very short or very long RRI). However, since HRnV is a novel representation of beat-to-beat variations in ECG, many technical issues need to be addressed in future research. For instance, as shown in Table 2, SampEn became larger when the available number of data points was less than 200 [19], suggesting that additional research is required to investigate its applicability to short ECG records. Moreover, parameters NN50*n* and pNN50*n* are newly introduced in HRnV representation only. They characterize the number of times that the absolute difference between two successive RR_*n*_I sequences exceeds 50×*n* ms, by assuming that the absolute difference may be magnified when the corresponding RR_*n*_I is *n* times longer than RRI. Thus, in-depth investigations are required in the selection of appropriate thresholds. More importantly, physiological interpretations of the HRnV parameters and their normal values [29] need to be determined through numerous research. One example is the identification of frequency bands that correlate with certain physiological phenomenon. In the current analysis, the conventional cut-off values were adopted (i.e., ≤0.04 Hz as very low frequency range; 0.04–0.15 Hz as low frequency range; 0.15–0.4 Hz as high frequency range). With the increase in *n*, frequency domain analysis may need to be changed accordingly.

Beyond its use in risk stratification of ED patients with chest pain, HRnV foresees potentials in many clinical domains where conventional HRV has been extensively investigated [46–49]. With the augmented RR_*n*_I and RR_*n*_I_*m*_ sequences, HRnV could possibly capture more dynamic changes in cardiac rhythms than HRV. This capability enables the extraction of extra information from limited raw ECGs. This study utilized HRnV parameters as independent risk factors and analyzed them with traditional biostatistical methods. There are multiple ways to use HRnV parameters, e.g., each set of HRnV parameters can be analyzed individually and are subsequently combined with an ensemble learning [50] (a special type of machine learning algorithm) architecture to reach a decision. However, artificial intelligence and machine learning methods generally create black-box predictive models, making interpretation a challenge [51].

### Limitations

This study has several limitations. First, no scoring tool was developed for practical clinical use. The primary aim of this study was to demonstrate the feasibility of using HRnV parameters and common risk factors to build predictive models. Second, the HRnV model was evaluated with LOOCV strategy due to small sample size. Ideally, separate patient cohorts are needed to train and test prediction models. When a new scoring tool is developed, it is necessary to conduct external validations on cohorts with diverse patient characteristics. Furthermore, properly designed clinical pathways are needed as well. Third, the patients included in this study were mainly from the high acuity group, resulting in a higher 30-day MACE rate (i.e., 31%) compared to other similar studies [10, 39]. As a result, the generalizability of the HRnV model developed in this study may be uncertain in other patient cohorts. Fourth, the calculated HRnV and HRV parameters depended on the choice of tools and methods for ECG signal analysis. Thus, the values of these parameters may vary across studies. Lastly, the physiology of HRnV and interpretations of its measures are mostly unknown; calculation of some parameters also needs to be standardized. All these require future collaborative efforts between clinicians and scientists to address.

## Conclusions

In this study, we proposed a novel HRnV representation and investigated the use of HRnV and established risk factors to develop a predictive model for risk stratification of patients with chest pain in the ED. Multiple HRnV parameters were found as statistically significant predictors, which effectively augmented conventional HRV, vital signs, troponin, and cardiac risk factors in building an effective model with good discrimination performance. The HRnV model outperformed the HEART, TIMI, and GRACE scores in the ROC analysis. It also demonstrated its capability in identifying low-risk patients, which may enable its use in building a new clinical pathway. Moving forward, we suggest further development of a point-based, ready-to-use HRnV risk stratification tool. Although some issues remain to be addressed, we hope to stimulate a new stream of research on HRnV. We believe that future endeavors in this field will lead to the possibility of in-depth evaluation of the associations between HRnV measures and various human diseases.

## List of abbreviations

ACS: acute coronary syndrome
ADP: accelerated diagnostic protocol
AMI: acute myocardial infarction
ANS: autonomic nervous system
AUC: area under the curve
ApEn: approximate entropy
CABG: coronary artery bypass graft
DFA: detrended fluctuation analysis
ED: emergency department
EHR: electronic health records
GRACE: Global Registry of Acute Coronary Events
HEART: History, ECG, Age, Risk factors and Troponin
HF: high frequency
HRnV: Heart Rate n-Variability
HRV: heart rate variability
LF: low frequency
LOOCV: leave-one-out cross-validation
MACE: major adverse cardiac events
Mean NN: average of R-R intervals
MI: myocardial infarction
NN50: the number of times that the absolute difference between 2 successive R-R intervals exceeds 50 ms
NN50n: the number of times that the absolute difference between 2 successive RRnI/RRnIm sequences exceeds 50×n ms
NPV: negative predictive value
PACS: Patient Acuity Category Scale
PCI: percutaneous coronary intervention
pNN50: NN50 divided by the total number of R-R intervals
pNN50n: NN50n divided by the total number of RRnI/RRnIm sequences
PPV: positive predictive value
RMSSD: square root of the mean squared differences between R-R intervals
ROC: receiver operating characteristic
RRI: R-R interval
SampEn: sample entropy
SD: standard deviation
SDNN: standard deviation of R-R intervals
STEMI: ST-elevation myocardial infarction
TIMI: Thrombolysis in Myocardial Infarction
VLF: very low frequency

## Declarations

## Ethics approval and consent to participate

The ethical approval was obtained from the Centralized Institutional Review Board (CIRB, Ref: 2014/584/C) of SingHealth, in which patient consent was waived.

## Consent for publication

Not applicable

## Availability of data and materials

The datasets used and/or analysed during the current study are available from the corresponding author on reasonable request.

## Authors’ contributions

NL invented the HRnV representation, conceived the study, supervised the project, and wrote the first draft of the manuscript. NL, DG, ZXK, and FX performed the analyses. All authors contributed to evaluation of the HRnV measures, interpretation of the results, and revision of the manuscript. All authors approved the final manuscript.

## Acknowledgements

We would like to thank and acknowledge the contributions of doctors, nurses, and clinical research coordinators from the Department of Emergency Medicine, Singapore General Hospital.

